# Co-administration of simvastatin mitigates clozapine-induced lifespan extension and mitochondrial functional recovery in *Drosophila melanogaster*

**DOI:** 10.64898/2026.10.19.760135

**Authors:** Katherine R.H Mortimer, Jacob Reed, Nicoleta Moisoi, Melissa Mather, Mohammad Zia Ul Haq Katshu, Reinhard Stöger, Lisa Chakrabarti

**Affiliations:** School of Veterinary Medicine and Science, University of Nottingham, Nottingham, United Kingdom; School of Pharmacy, De Montfort University, Leicester, United Kingdom; Optics and Photonics Group, Faculty of Engineering, University of Nottingham, UK; Institute of Mental Health, School of Medicine, University of Nottingham, Nottingham, United Kingdom; Nottinghamshire Healthcare NHS Foundation Trust, Nottingham, United Kingdom; School of Biosciences, University of Nottingham, Nottingham, United Kingdom

## Abstract

Antipsychotics and statins can be co-prescribed to combat the increased risk of high blood cholesterol with antipsychotic use. However, statins have been shown to interfere with mitochondrial functioning through reduced coenzyme Q expression, and second-generation antipsychotic, clozapine, has also demonstrated induced mitochondrial respiratory deficits in cell models. Despite this convergence, the combined effect of statins and antipsychotics on mitochondrial function, and mechanisms of overall health, remain poorly understood. Here we exploited the cholesterol auxotrophy of *Drosophila melanogaster* to investigate the effects of clozapine, simvastatin and their combination on lifespan and mitochondrial bioenergetics independently of cholesterol biosynthesis. Clozapine alone significantly extended median survival, an effect attenuated by simvastatin co-treatment. High resolution respirometry showed signs of clozapine-induced mitochondrial differences diminished with age, but this adaptation was also attenuated by simvastatin co-treatment. Using a spin probe (TEMPOL) with nitrogen-vacancy diamond quantum sensing, we directly measured free radical production and found that clozapine and simvastatin significantly increased free radical generation relative to the untreated control in early life stages. We also observed preliminary evidence of sex-specific differences in treatment response. Together, these findings suggest that clozapine induces an age-related mitochondrial adaptive response that is associated with extended lifespan and mitigated by simvastatin treatment.

## 2. Introduction

The antipsychotic medication clozapine, while highly efficacious at treating psychosis, is associated with many side-effects ^1,2^. The metabolic side-effects of clozapine use can trigger metabolic syndrome (MetS) through rapid weight gain, elevated blood glucose and cholesterol levels, and eventual development of type II diabetes and obesity ^3–6^. These side-effects are particularly concerning when considering the already elevated risk of cardiovascular disease associated with schizophrenia ^3^. Statins are commonly prescribed to treat high blood cholesterol and the incidence of statin prescription in individuals aged 30-59 years with schizophrenia is 2-3-fold higher compared to those without ^7^. Statins act on the rate-limiting step of the mevalonate pathway by inhibiting 3-hydroxy-3-methylglutaryl-CoA reductase (HMGCR) ^8^. The mevalonate pathway is responsible for the biosynthesis of cholesterol and so is a prime target for managing hypercholesterolemia. Statins have been shown to reduce total cholesterol, LDL and triglyceride levels after one month of treatment in males already taking antipsychotics ^9^. However, it has also been shown that statins failed to reduce circulating LDL levels in patients taking second-generation antipsychotics ^10^.

In mammals, clozapine upregulates sterol regulatory element-binding proteins (SREBPs) ^11,12^. SREBPs are transcriptional regulators of mevalonate pathway and fatty acid synthesis enzymes, with consequent increases in cholesterol and free fatty acid levels linked to mitochondrial dysfunction and hypercholesterolemia, which contribute to MetS. The fruit fly, *Drosophila melanogaster*, is a commonly used model for studying human metabolism with many metabolic pathways conserved between flies and humans^13^. *D. melanogaster* is used to model obesity and mitochondrial dysfunction in Parkinsons’s disease, as well as for testing of therapeutics ^13^. Although *D. melanogaster* do not have livers, their fat bodies perform many analogous functions including triglyceride and glycogen storage and nutrient sensing. The *D. melanogaster* mevalonate pathway lacks the cholesterol synthesis branch, meaning flies must obtain cholesterol exclusively from dietary sources ^14,15^. This represents an interesting scenario for research. By eliminating cholesterol as a pathway output, we have the opportunity to independently interrogate the downstream consequences of mevalonate pathway upregulation. In *D. melanogaster*, the mevalonate pathway drives production of non-sterol isoprenoids, including coenzyme Q (CoQ) and prenylated proteins ^16^, enabling us to dissect the contribution of these outputs to clozapine-associated metabolic dysfunction independent of cholesterol. Previously, simvastatin was shown to extend lifespan through decreased protein prenylation ^17^, indicating that mevalonate pathway-derived isoprenoids are sufficient to influence health span in *D. melanogaster*.

The mevalonate pathway is closely tied to mitochondrial function through production of CoQ which is an essential electron carrier in the mitochondrial respiratory chain ^18^. Both clozapine and simvastatin interfere with normal mitochondrial functioning ^19–22^. Clozapine altered mitochondrial morphology and reduced ATP production in multiple cell models ^19,20^. Simvastatin also reduced ATP production in human liver cells, through reduced levels of mitochondrial coenzyme Q10 (CoQ10) ^22^. Based on these findings, we observe the effect of co-administration of these medications on mitochondrial functioning in a whole organism to reflect the treatment regime in clinic.

Our study utilizes a multi-method approach to determine the effects of simvastatin and clozapine co-treatment on life span, climbing ability, mitochondrial physiology and free radical production in *D. melanogaster.* We also performed a CApillary FEeder (CAFE) assay to measure appetite and observe any potential dietary preferences between treatment groups ^23,24^. Initial studies on mitochondrial physiology and dietary preferences were performed on both males and females to determine any sex differences.

## 3. Methods

### D. melanogaster husbandry and drug treatment

Fly stocks were maintained on standard cornmeal media (Nutri-Fly BF, FLY1034) at 25° C in a 12:12 light-dark cycle. The experiments were performed on wildtype (genotype w1118). Clozapine and simvastatin were dissolved in DMSO to a 100 mM stock solution and stored at -20 °C until use. They were then dissolved in H_2_O to the desired concentration and 200 μL was mixed with 100 mg of dry yeast to form yeast paste ^25,26^.

### Life span and climbing assay

At day 10, flies were added to a new vial of food with the addition of yeast paste containing 1 mM clozapine, 1 mM clozapine + 0.24 mM of simvastatin, 0.24 mM of simvastatin or DMSO as a control. The dose used was based on previous studies ^17,27^. Three vials were used for each treatment group containing 15-20 flies each. Fly food and yeast paste were changed every 3 days and the number of live flies recorded each day. Data are presented as Kaplan–

Meier survival distributions and significance determined by Log-rank tests. A climbing assay was conducted prior to drug treatment and after 21, 42 and 63 days of treatment. Flies were transferred to the climbing assay apparatus, knocked to the bottom of the climbing vial and allowed 18 s to climb 5 cm up the vial. The percentage of flies passing the 5 cm mark was recorded and the experiment repeated 3-5 times ^28^.

### High resolution respirometry

Mitochondrial respiratory function was assessed using an Oxygraph-2k (O2k) high-resolution FluoRespirometer (Oroboros Instruments, AT). Respirometry data were collected at 25°C in MiR05 buffer with a stirring speed of 750 r.p.m. Before the experiments began, an air calibration was performed. Female and male flies were anaesthetized on ice after 14 days of treatment, and for the males only experiment after 45 days of treatment. 3 whole flies from each treatment group were homogenized in 300μL of Mir05. 200μL of homogenate was added to each chamber of the respirometer. A SUIT protocol (SUIT-005 O2 pfi D11) was used to measure oxygen consumption through the F pathway (where fatty acids feed electrons to FADH_2_) and N pathway (NADH) and S pathway (succinate-linked respiration) ^29^. The F pathway is measured by adding malate (0.1mM) and octanoylcartinine (0.5mM) to the chambers. This provides a reading of oxygen consumption through stimulation of fatty acid oxidation (FAO) in Leak phase. ADP (2.5mM) is added to activate OXPHOS. Then complex I (CI) substrates for the N pathway, Pyruvate (5mM) and Proline (10mM) are added. Then succinate (10mM) is added to activate complex II (CII) and finally CI and CI + CII inhibitors rotenone (0.25uM) and antimycin A (2.5uM). This protocol gave us oxygen consumption readings for F(Leak), F(OXPHOS), CI OXPHOS and CI + CII OXPHOS.

### CApillary FEeder (CAFE) assay

To analyze food preferences, the CAFE method was utilized following a design used in previous studies ^23,24^. A stock solution of clozapine and simvastatin (100 mM) were dissolved in 5% sucrose solution to a concentration of 1mM and 0.24mM respectively. 5μL capillaries were filled with clozapine sucrose solution, simvastatin sucrose solution, clozapine + simvastatin sucrose solution, or sucrose solution with corresponding volumes of DMSO as control. Female and male flies were used in this study. 6 flies were added to a CAFE apparatus with 5x 5μL capillaries. The microcapillaries were changed daily, and the volume remaining was recorded. The (volume left in the capillaries – volume evaporated)/number of flies was used to determine the volume of solution consumed per fly. Flies were exposed to microcapillaries for 72 hours. The study was repeated 3 times.

### Quantum Sensing of Free Radicals Production

Free radicals were quantified using a quantum sensing approach based on the spin-dependent photoluminescence (PL) of nitrogen-vacancy (NV) centres in nanodiamonds^30^. An applied magnetic field modulates the NV spin state, producing a PL contrast that reports on nearby paramagnetic species. Selectivity was achieved using the spin probe TEMPOL, a superoxide dismutase 2 (SOD2) mimic that quenches PL in its paramagnetic form but is converted to a diamagnetic derivative upon scavenging hydroxyl and superoxide radicals, restoring PL contrast in proportion to the radicals scavenged.

Male flies were exposed to 1mM clozapine, 1mM clozapine + 0.24 mM of simvastatin, 0.24mM of simvastatin or DMSO in yeast paste for 14 days. Three flies were pooled into an 1.5ml tube (4 biological repeats per treatment group) and homogenised directly into 100μL of TEMPOL (0.0025 M). The homogenates were frozen at -20° C. Carboxylated fluorescent nanodiamonds (FND-Biotech, Taiwan) were drop-cast onto glass coverslips and imaged on an inverted fluorescence microscope (Olympus IX83) fitted with a 60X oil-immersion objective (NA = 1.49). NV centres were excited at 550 nm (CoolLED pE-4000, 545 ± 15nm bandpass) and PL was collected through a 575nm longpass filter onto a sCMOS camera (Photometrics Prime 95B), controlled via Olympus CellSens. A 40 ± 1mT magnetic field, square-wave modulated at 0.5 Hz, was applied using an electromagnet driven by a signal generator, with PL recorded over 4 cycles (210 frames, 30ms exposure). PL contrast was calculated in ImageJ as the normalized difference in average PL between ‘field-on’ and ‘field-off’ states across selected regions of interest.

### Statistical analysis

For respirometry, climbing assay and CAFE assay, data are expressed as mean ± SEM. To calculate statistical significance (*P* < .05), a one-way or two-way ANOVA with Šidák correction or Tukey’s test for multiple comparisons was performed where appropriate. For the climbing assay and CAFE assay, a two-way ANOVA was used to determine the significance of changes within each group over time, as well as between groups at each time point. For respirometry, two-way ANOVAs were used as measurements were taken from the same biological replicates over time. Sample size is indicated in figure legends. For the NV⁻ diamond contrast data, treatment values were normalized to each diamond’s own DMSO control and compared to the null value of 1 using one-sample t-tests (Holm-Bonferroni corrected), while differences between treatments were assessed using a repeated-measures one-way ANOVA followed by paired t-tests (Holm-Bonferroni corrected). Asterisks indicate as follows: **** *P* < .0001 *** *P* < .001, ** P < .01, * *P* < .05. Kaplan-Meier survival curves were constructed for each treatment group, and differences in survival distributions were assessed using the global log-rank test. Pairwise log-rank tests were conducted between all treatment combinations, with p-values adjusted for multiple comparisons using the Benjamini-Hochberg method. To characterize the underlying mortality dynamics, a Gompertz proportional hazards model was fitted to the survival data, enabling estimation of the aging rate parameter (β) and baseline mortality (α) for each group. Statistical analyses and graphs were made in R (version 4.4.0).

## 4. Results

### Clozapine increased lifespan and slowed ageing rate in wildtype male flies

Flies exposed to 1 mM clozapine in yeast paste supplemented with their food showed a significantly increased lifespan (Fig. 1A) and reduced age-dependent mortality (Fig. 1B-C). The median survival for clozapine exposed flies was 92 days, compared to 73 for DMSO treated control flies. Co-treatment with simvastatin significantly attenuated this increase in lifespan, with flies exposed to both clozapine and simvastatin having a median survival of 73 days (Table 1). Simvastatin alone did not significantly alter lifespan.

Mortality modeling indicated that clozapine treatment reduced the age-dependent increase in mortality (Table 2). In contrast, co-treatment of clozapine and simvastatin reduced baseline mortality but increased age-dependent mortality, leading to non-significant alterations in lifespan.

**Figure 1:**
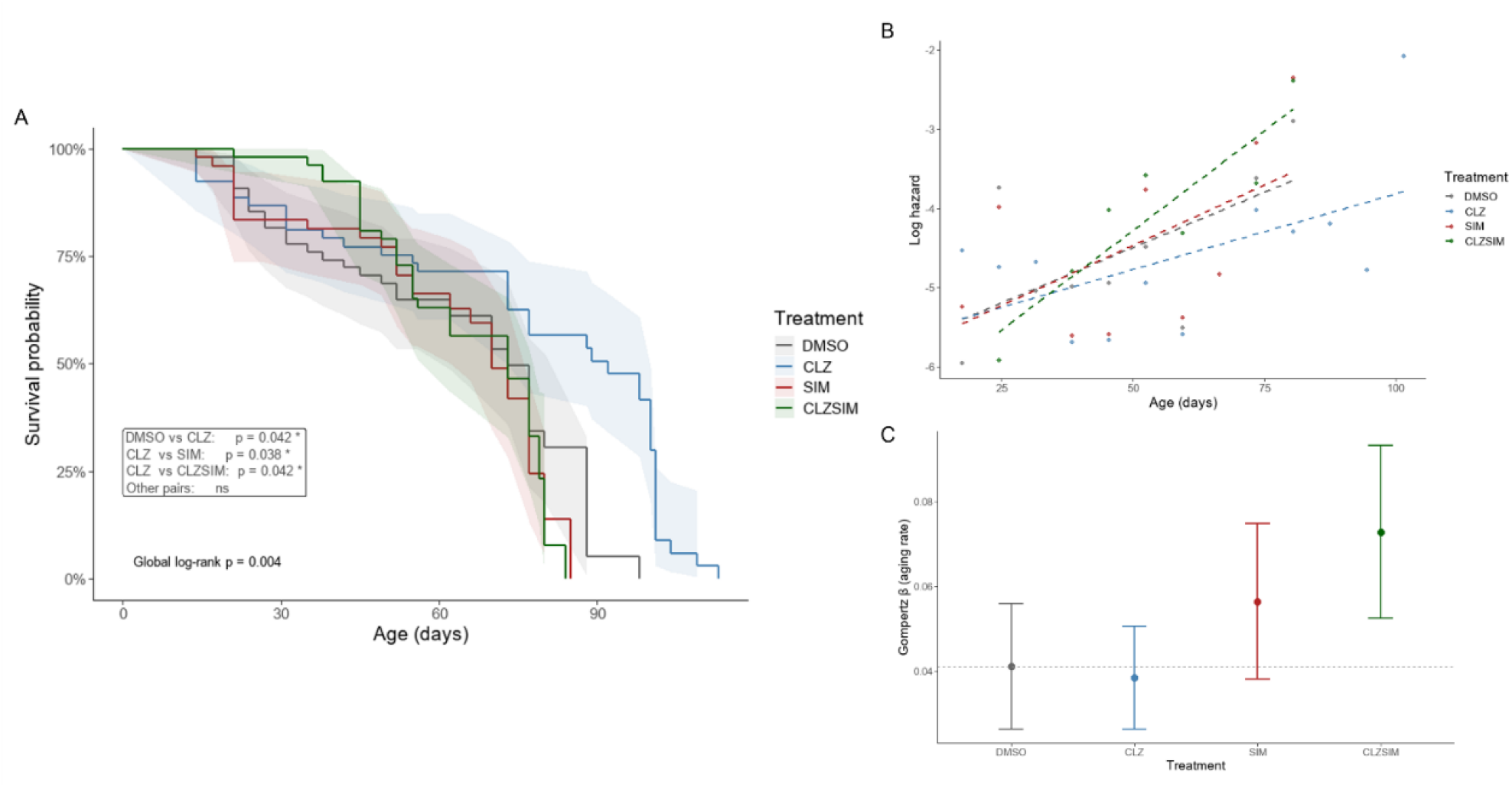
Survival analysis of male *Drosophila melanogaster* exposed to DMSO (vehicle control), clozapine (CLZ), simvastatin (SIM), or clozapine and simvastatin combined (CLZSIM). Kaplan-Meier survival curves with 95% confidence intervals (shaded), n=55 per treatment group. Significant pairwise comparisons (BH-adjusted log-rank test) are indicated **(A)**. Log-transformed empirical hazard rate plotted against age, with linear regression lines (dashed) fitted per group. Parallel lines indicate equivalent aging rates (β); vertical offset indicates differences in baseline mortality (α) **(B)**. Gompertz aging rate parameter (β) estimated per treatment group from independently fitted Gompertz proportional hazards models. Points represent maximum likelihood estimates; error bars represent 95% confidence intervals. Significance relative to DMSO was assessed using a combined Gompertz proportional hazards model (CLZ: * p < 0.05; SIM and CLZSIM: ns). Dashed line indicates DMSO reference value **(C).**

**Table 1:** Median survival estimated from Kaplan-Meier survival analysis for each treatment group. 95% confidence intervals are calculated using the log-log transformation method. DMSO (vehicle control), clozapine (CLZ), simvastatin (SIM), and clozapine and simvastatin combined (CLZSIM).

| Treatment | Median survival (days) | 95% CI |
| --- | --- | --- |
| DMSO | 73 | 62–88 |
| CLZ | 92 | 73–100 |
| SIM | 70 | 66–77 |
| CLZSIM | 73 | 62–79 |

**Table 2:** Gompertz aging rate (β) per treatment group estimated from independently fitted models. DMSO (vehicle control), clozapine (CLZ), simvastatin (SIM), and clozapine and simvastatin combined (CLZSIM). Significance vs DMSO assessed using a combined Gompertz proportional hazards model (CLZ: * p < 0.05; SIM and CLZSIM: ns)

| Treatment | $\beta$ (aging rate) | 95% CI | $\alpha$ (baseline mortality) | 95% CI |
| --- | --- | --- | --- | --- |
| DMSO | 0.041 | 0.026–0.056 | 0.00202 | 0.00086–0.00473 |
| CLZ* | 0.038 | 0.026–0.051 | 0.00135 | 0.00055–0.00332 |
| SIM | 0.056 | 0.038–0.075 | 0.00101 | 0.00034–0.00295 |
| CLZSIM | 0.073 | 0.052–0.093 | 0.00037 | 0.00011–0.00131 |

### Male and female flies showed no differences in food preference between treatment groups

To ensure that any effects observed between treatment groups were not due to the drug causing food aversion, a CAFE assay was conducted. Male and female flies were exposed to microcapillaries containing 5% sucrose supplemented with clozapine and/or simvastatin and a vehicle control. After 72 hours, there were no statistically significant differences in food intake between groups in the flies (Fig. 2).

**Figure 2:**
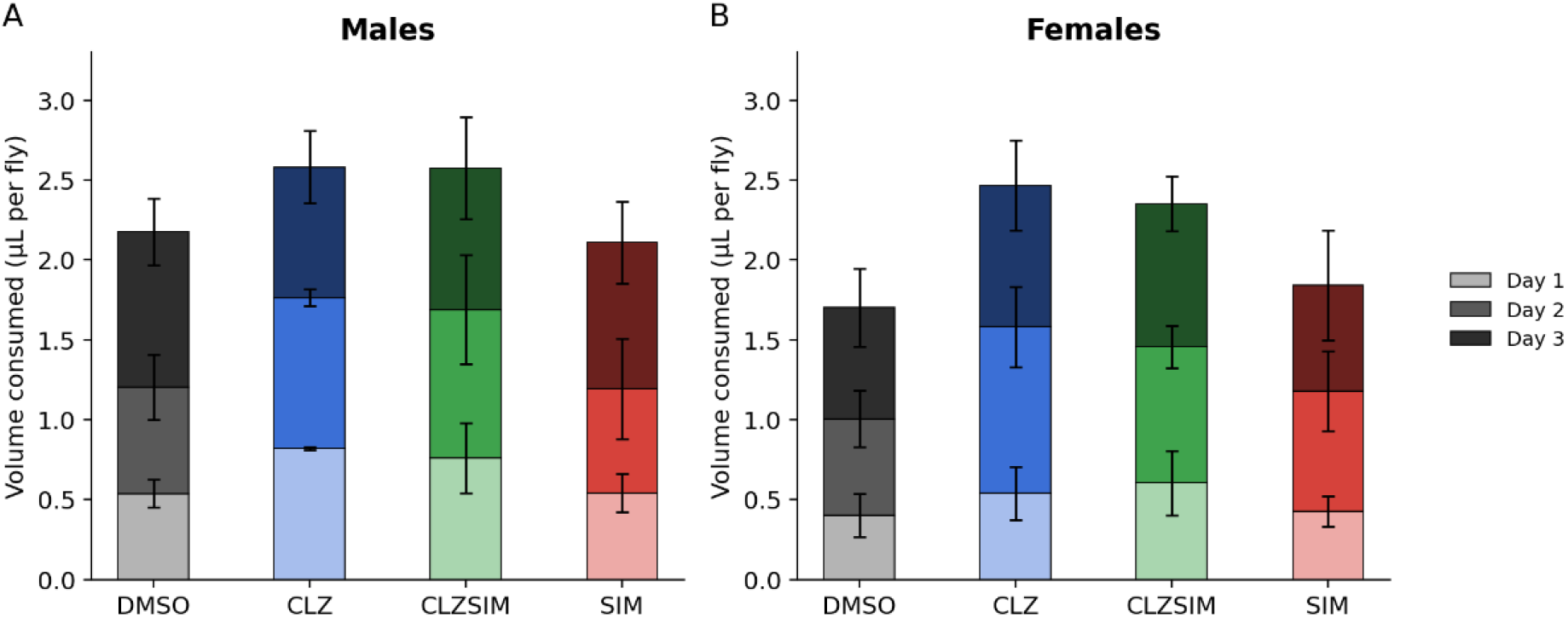
Food preferences in male and female *D. melanogaster.* A CAFE assay to assess food preference and appetite in flies exposed to DMSO (vehicle control), clozapine (CLZ), simvastatin (SIM), or clozapine and simvastatin combined (CLZSIM) in 5% sucrose solution. N=3, n=5-7 flies. Error bars represent ± SEM.

### Early signs of mitochondrial changes with clozapine exposure diminished with age

After 14 days of treatment, male flies treated with clozapine (both alone and with simvastatin) showed mitochondrial respirometry differences through complex I and complex I + II OXPHOS (Fig. 3B). With 45 days of treatment, only the co-treated male flies demonstrated continued respirometry changes (Fig. 3D). Additionally, with age, clozapine-treated, simvastatin-treated and co-treated flies showed increased respiration through FAO compared to N linked respiration (CI). At day 45, this increase was significant in both groups exposed to clozapine compared to DMSO control flies (Fig. 3E).

**Figure 3:**
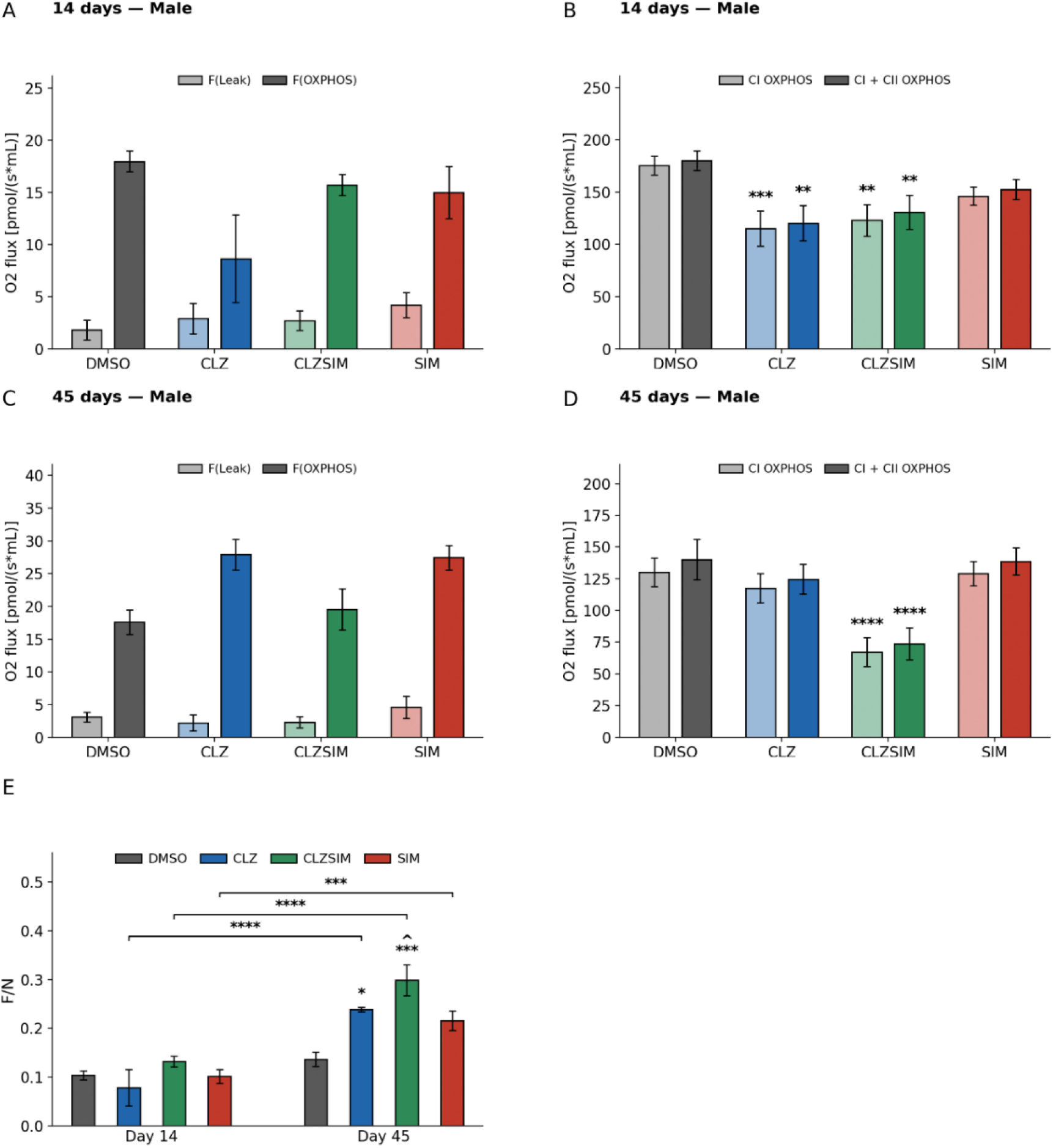
Respirometry analysis of male *D. melanogaster* exposed to DMSO (vehicle control), clozapine (CLZ), simvastatin (SIM), or clozapine and simvastatin combined (CLZSIM). Oxygen flux values per group obtained after 14 days in F(Leak) and F(OXPHOS) state. Values represent a pool of 3 flies per run **(A)**. Oxygen flux values per group obtained after 14 days in CI(OXPHOS) and CI + CII (OXPHOS), CLZ oxygen flux values were significantly lower than the control in CI(OXPHOS) (*P* = .0009, 95% C.I. = [21.33, 98.99]) and CI + CII(OXPHOS) (*P* = .001, 95% C.I. = [20.92, 98.58]) as were flux values in the CLZSIM group in CI (*P* = .002, 95% C.I. = [16.02, 88.66]) and CI + CII(OXPHOS) (*P* = .004, 95% C.I. = [13.02, 85.66]) **(B)**. Oxygen flux values per group obtained after 45 days in F(Leak) and F(OXPHOS) state **(C)**. Oxygen flux values obtained after 45 days in CI(OXPHOS) and CI + CII (OXPHOS), CLZSIM oxygen flux values were significantly lower compared to controls in OXPHOS through CI (*P* < .0001, 95% C.I. = [30.37, 95.40]) and CI + CII (*P* <.0001, 95% C.I. = [33.25, 99.19]) **(D)**. Ratio of oxygen flux through F(OXPHOS) (F) compared to CI(OXPHOS) (N) between 14 and 45 days was significantly increased in CLZ (*P* <.0001, 95% C.I. = [-0.22, -0.10]), CLZSIM (*P* <.0001, 95% C.I. = [-0.22, -0.12]) and SIM (*P* =.00058, 95% C.I. = [-0.172, -0.056]) groups. At day 45, F/N rati for CLZ (*P* = .02, 95% C.I. = [-0.19, -0.011]) and CLZSIM (*P* = .0002, 95% C.I. = [-0.25, - 0.078]) were significantly increased compared to DMSO control **(E)**. Error bars represent ± SEM. * Compared to control, ^ compared to SIM.

### No significant changes in mitochondrial function were observed in female Drosophila melanogaster after 14 days of treatment

In contrast to male flies, female flies showed no respirometry differences between treatment groups following 14 days of treatment with clozapine and/or simvastatin compared to the control in either F-linked or CI/CI + CII linked respiration (Fig. 4).

**Figure 4:**
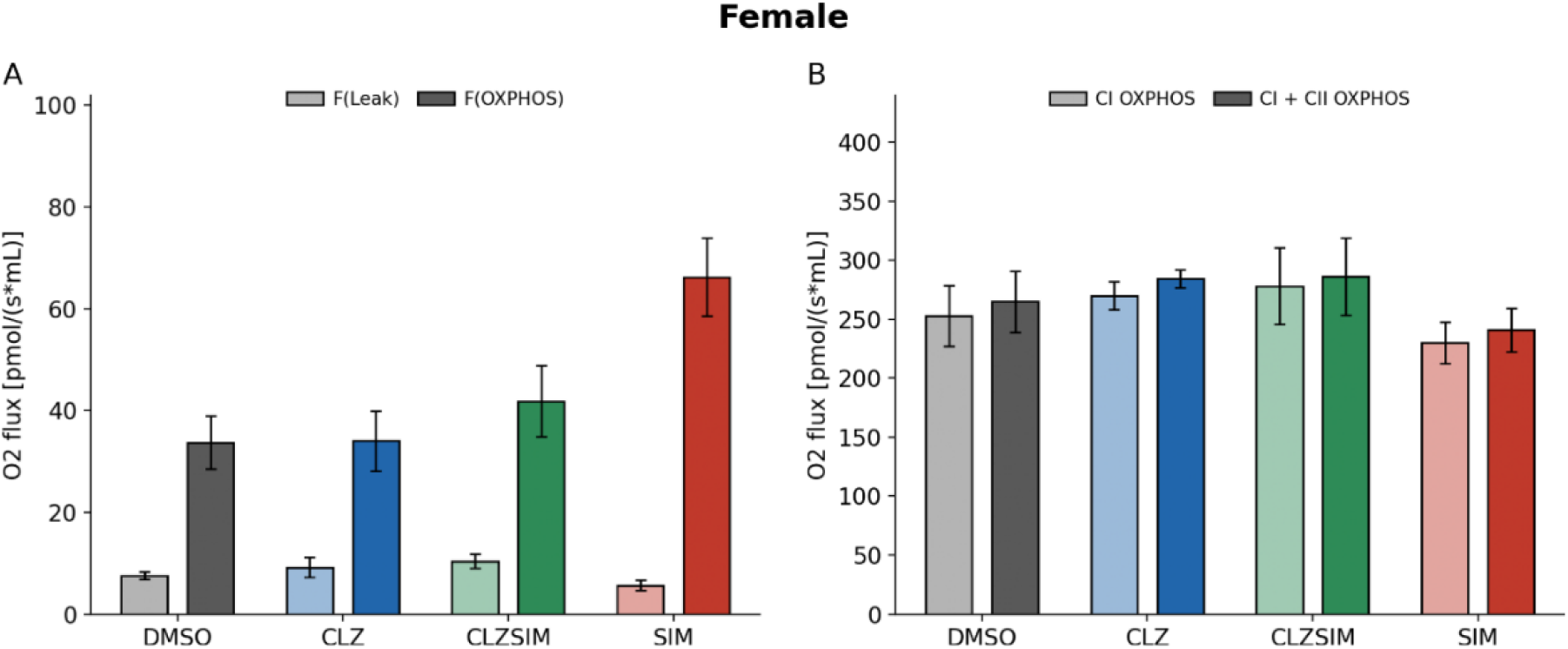
Respirometry analysis of female *D. melanogaster* exposed to DMSO (vehicle control), clozapine (CLZ), simvastatin (SIM), or clozapine and simvastatin combined (CLZSIM). Oxygen flux values per group obtained after 14 days in F(Leak) and F(OXPHOS) state. Values represent a pool of 3 flies per run **(A)**. Oxygen flux values per group obtained after 14 days in CI(OXPHOS) and CI + CII (OXPHOS) **(B)**. Error bars represent ± SEM.

### Quantity of free radicals is increased in all treatment groups compared to the control

Quantum sensing of free radicals using NV nanodiamonds was conducted on male fly samples after 14 days of exposure to clozapine, simvastatin, both, or a DMSO control (Fig. 5A). All treatment groups showed significantly increased quantity of free radicals compared to the control. Clozapine exposed flies showed the highest quantity of free radicals, followed by the co-treatment, then simvastatin.

### Clozapine improved climbing ability after 42 days of treatment but did not completely mitigate age-related climbing deficits

The climbing ability of the flies diminished with age. A two-way ANOVA revealed a significant Treatment × Day interaction (F (9,52) =3.17, P = .004), indicating that treatment effects on climbing ability varied over time; treatments were therefore compared separately at each timepoint. After 42 days, the clozapine-treated group showed increased climbing ability compared to the control group (Fig. 5B). This increase was no longer present after 63 days of treatment. At 63 days, clozapine and simvastatin co-treatment showed significant deficits in climbing ability compared to all other groups.

**Figure 5:**
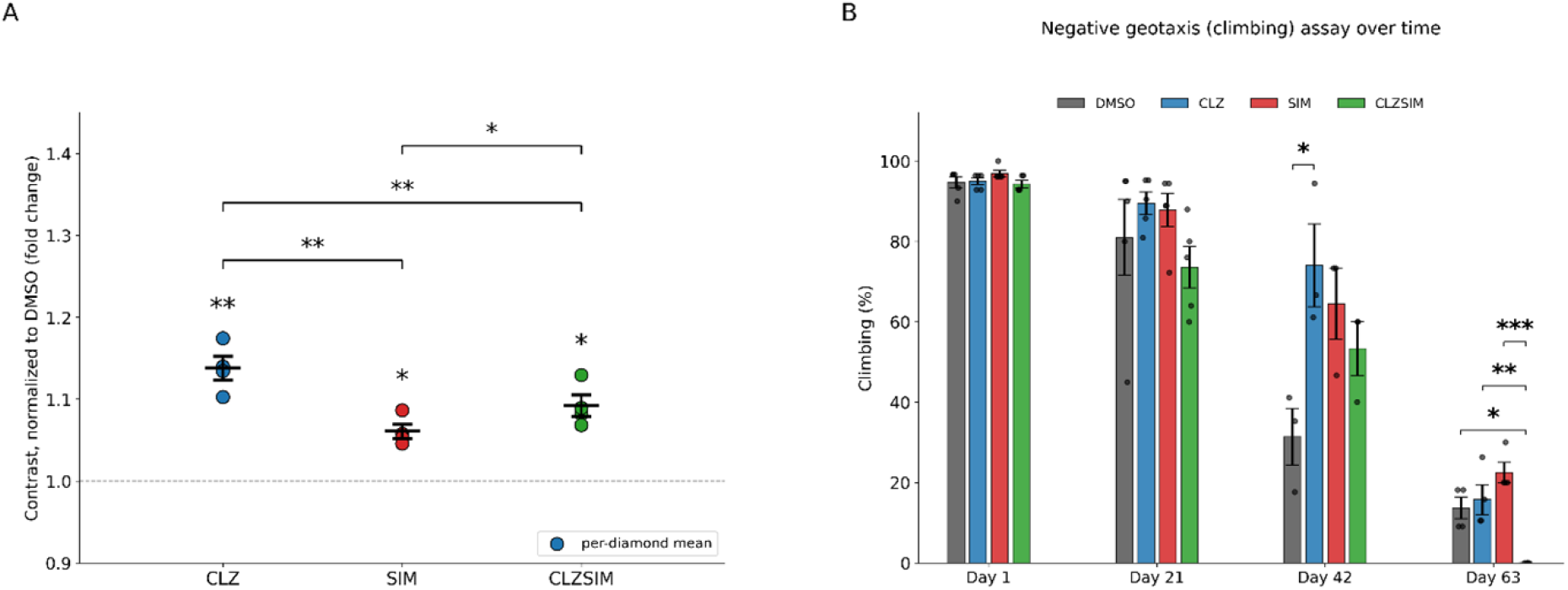
Free radical production analysis and climbing assay in male *D. melanogaster* treated with clozapine (CLZ), simvastatin (SIM), co-treatment (CLZSIM) or DMSO control. NV⁻ contrast was measured across four NV diamonds, with each treatment normalized to its own DMSO control (dashed line, =1). CLZ (P = .0077), SIM (P = .0120), and CLZSIM-exposed diamonds (P = .0120) all showed significantly increased contrast compared to DMSO (Holm-corrected one-sample t-tests). Repeated measures one-way ANOVA revealed a significant effect of treatment (F(2,6), P = .0001). Pairwise comparisons (paired t-tests, Holm-corrected) showed CLZ had significantly greater contrast than SIM (P = .008) and CLZSIM (P = .008), and CLZSIM was significantly greater than SIM (P = .0220) **(A)**. Climbing ability assessed on day 1, 21, 42 and 63. At day 42, CLZ-exposed flies had significantly improved climbing ability compared to control flies (P = .0283, 95% C.I. = [−80.58, −4.83]). At day 63, CLZSIM-exposed flies showed significantly reduced climbing ability compared to control (P = .0136, 95% C.I. = [−24.54, −2.73]), CLZ (P = .0049, 95% C.I. = [4.88, 26.70]), and SIM (P = .0003, 95% C.I. = [−33.41, −11.59]) **(B)**.

## 5. Discussion

We found clozapine-treated *D. melanogaster* had a significantly extended lifespan and reduced ageing rate. Clozapine treatment (including clozapine and simvastatin co-treatment) also induced mitochondrial respiratory chain dysfunction through CI and CI + CII in early life, but after 45 days of treatment, only the co-treatment group showed continued dysfunction. In combination, these results show that clozapine induces an adaptive response over time that not only mitigates the mitochondrial changes, but also increases lifespan significantly compared to controls. Since the co-treatment group showed no recovery of mitochondrial function after 45 days and no extended lifespan, this suggests that the adaptive response by clozapine is attenuated by simvastatin. From our data, we see a distinct increase in FAO-linked respiration over time, which is particularly prominent in clozapine-treated groups. Clozapine has been known to upregulate many genes involved in lipogenesis including SREBP target genes ^31^. In *D. melanogaster,* SREBP exclusively upregulates fatty acid synthesis ^32^. Early life upregulation of fatty acid synthesis by clozapine may induce progressive overload of intracellular fatty acids, leading to accumulation of lipid droplets. This would elevate the substrate pool for FAO and may explain the increase in FAO-linked respiration at day 45. In dietary restriction models, an increase in fatty acid synthesis and FAO are associated with increased longevity in *D. melanogaster*, as inhibition of these pathways attenuates the lifespan-extending effect of dietary restriction ^33^. However, since this effect was seen in both clozapine-treated groups, it is unlikely to account for the increased longevity with clozapine alone.

At day 14, clozapine reduced CI activity. Lifespan extension by ETC dysfunction has been observed in *D. melanogaster* previously ^34,35^. Downregulation of genes encoding CI by RNAi was associated with significantly increased longevity ^34^. The exact mechanism for this lifespan extension is not known, but the possibility that it acts through reconfiguring metabolic gene expression to favour longevity aligns with our findings. Initially CI inhibition by clozapine led to an adaptive response that increased longevity and ameliorated mitochondrial ETC changes.

Clozapine has been shown to induce oxidative stress in rats and cell models ^36,37^. CI inhibition enhances ROS production, and the resulting oxidative stress further impairs CI activity ^38^, suggesting that clozapine triggers a positive feedback loop of ROS generation and CI inhibition. Consistent with this, superoxide levels, quantified using nitrogen-vacancy (NV) centre quantum diamond sensors, directly confirmed that after a 14-day exposure, clozapine significantly increased free-radical-associated photoluminescence contrast relative to DMSO controls, and this increase was significantly greater than that produced by simvastatin or co-treatment. This elevated free radical production may contribute to the reduced CI activity observed with clozapine treatment. Given that ROS increase with ageing yet can extend *D. melanogaster* lifespan through CI reverse electron transport (RET), in which electrons flow from reduced coenzyme Q (CoQ) back to CI ^39^, it may also form part of the adaptive longevity response underlying clozapine’s lifespan-extending effect. Simvastatin alone also produced a significant, although smaller, increase in free radical contrast relative to DMSO, while co-treatment flies showed an intermediate level of free radical production. As simvastatin can deplete CoQ, this partial attenuation of the clozapine-associated free radical signal in co-treated flies is consistent with reduced RET-derived ROS. This suggests a mechanistic explanation behind the loss of clozapine’s lifespan-extending effect when co-administered with simvastatin.

The effect of simvastatin on CoQ that has been shown to induce mitochondrial dysfunction in other studies is not reflected in our results ^22,40^. Simvastatin alone did not interfere with mitochondrial respiratory activity. This puts further emphasis on the combined effect of clozapine and simvastatin and the antagonist nature of the two medications. Spindler *et al.* (2012) reported that simvastatin significantly increased lifespan in *D. melanogaster* and attributed this effect primarily to reduced protein prenylation rather than alterations in CoQ levels ^17^. However, this study assessed CoQ levels using an LC-MS/MS method calibrated against CoQ10 standards. As *D. melanogaster* predominantly synthesizes CoQ9 rather than CoQ10, this approach may not have accurately captured changes in the functionally relevant ubiquinone isoform ^16^. In addition, we did not observe any changes in lifespan following simvastatin treatment. This discrepancy may partly reflect differences in treatment initiation: we began treatment at day 10 to ensure reproductive maturity and metabolic stabilization, whereas Spindler *et al.* initiated treatment at eclosion.

The decline in climbing ability observed in control flies between days 21 and 42 coincides with the period of peak mortality in that group, suggesting that the functional decline reflects age-related deterioration. Clozapine-treated flies showed significantly improved climbing ability relative to controls at day 42, while simvastatin- and co-treated flies showed a similar, non-significant trend. By day 63, this clozapine-associated improvement was no longer evident, indicating that this change was transient. By approximately day 60, the survival curves of simvastatin and co-treatment flies converge with the control; however, climbing data at day 63 reveal that co-treated (CLZSIM) flies had significantly poorer climbing ability than control, clozapine-, and simvastatin-treated flies, indicating that the combination of clozapine and simvastatin imposes an additional, disproportionate burden on late-life locomotor function beyond what would be predicted from survival alone. The lifespan extension conferred by clozapine did not coincide with a proportional prolongation of climbing ability, suggesting these phenotypes may be regulated through partially dissociable mechanisms. This is consistent with Wilson et al. (2020), who demonstrated that lifespan and age-related locomotor decline in *D. melanogaster* are modulated by distinct genetic factors ^41^.

We wished to address whether there were sex differences in the responses we saw in *D. melanogaster*. At 14 days of treatment, there were no observed significant differences in oxygen flux in any respiratory state in female flies. Female flies have been shown to be more resistant to oxidative stress than males and have a higher antioxidant capacity which might explain their resistance to alterations in mitochondrial function, following clozapine exposure

In this study on *D. melanogaster*, the increased lifespan and adaptive response to oxidative stress by clozapine is observed independent of cholesterol syntheis. Hypercholesterolemia in patients is a serious risk factor for cardiovascular disease and mitochondrial cholesterol accumulation is associated with impairments in OXPHOS ^43^.

Our findings are specific to simvastatin, a lipophilic statin with CoQ-lowering properties. Whether hydrophilic statins with reduced effect on CoQ production, such as pravastatin, interact similarly with clozapine’s adaptive response is unknown but would be highly relevant to statin selection in clinical practice. Once tested, if they do not yield a similar effect to simvastatin regarding mitochondrial physiology and mitigation of lifespan extension, such statins may present a preferable choice for patients requiring lipid-lowering therapy alongside clozapine.

## 6. Conclusion

This study is the first to observe the effects of co-administration of clozapine and simvastatin on mitochondrial function and lifespan in *D. melanogaster*. Our findings revealed a cholesterol-independent mechanism of clozapine that induced adaptive longevity through early life mitochondrial physiology effects and an increase in free radical production. This adaptive response was blunted by simvastatin treatment, suggesting a role for mevalonate pathway products in the life-span extension effects of clozapine.

## Supplementary Information

**Supplementary Figure 1:**
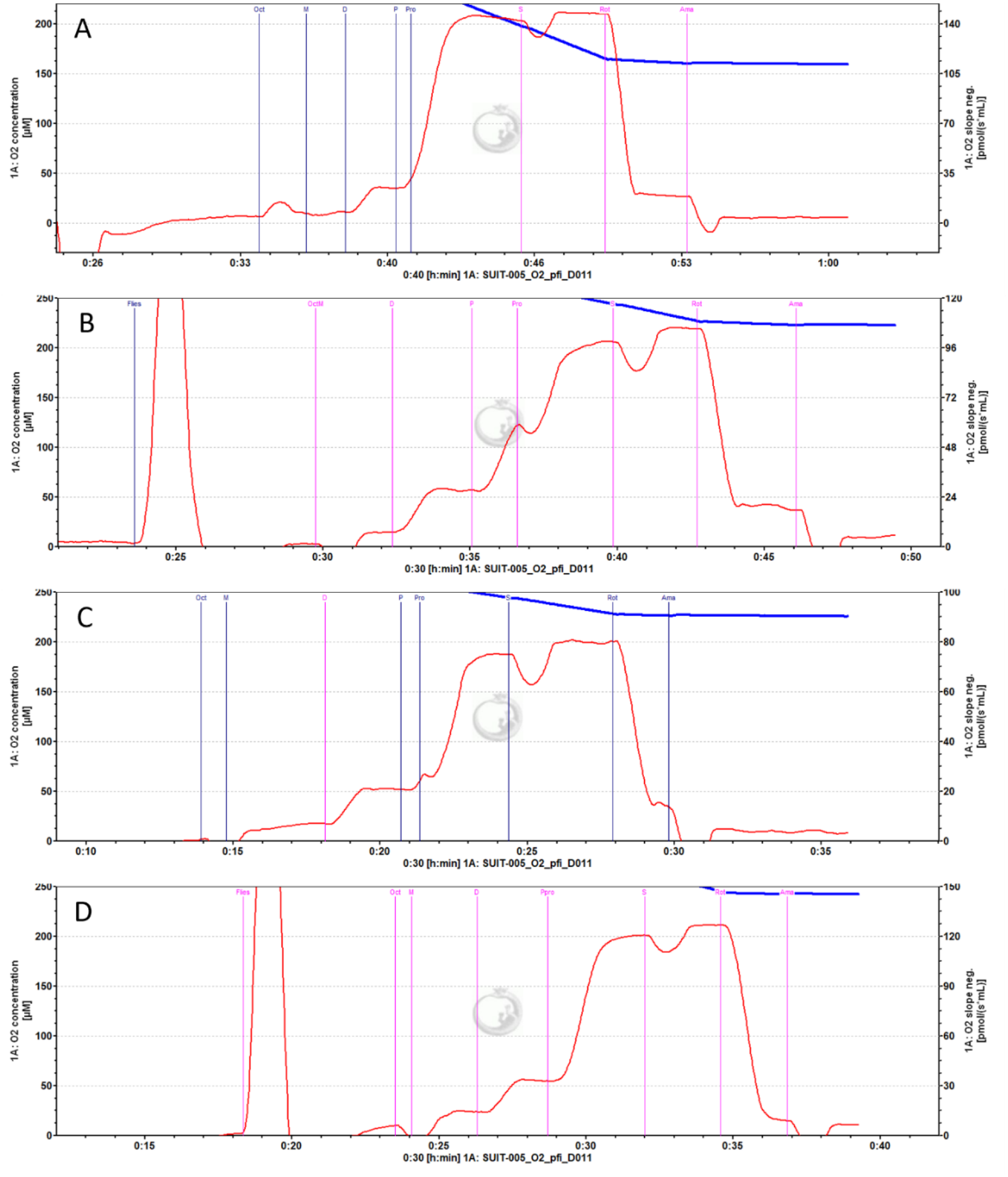
Example Datlab traces from male *D*. *melanogaster* exposed to **(A)** DMSO, **(B)** 1 mM clozapine, **(C)** 1 mM clozapine + 0.24 mM simvastatin **(D)** 0.24 mM simvastatin for 45 days. Oxygen concentration is shown in blue and oxygen flux in red. Protocol used was SUIT-005 D011 without uncoupler. Marks were made after stabilisation of oxygen flux for a few minutes following reagent administration.

**Supplementary Figure 2:**
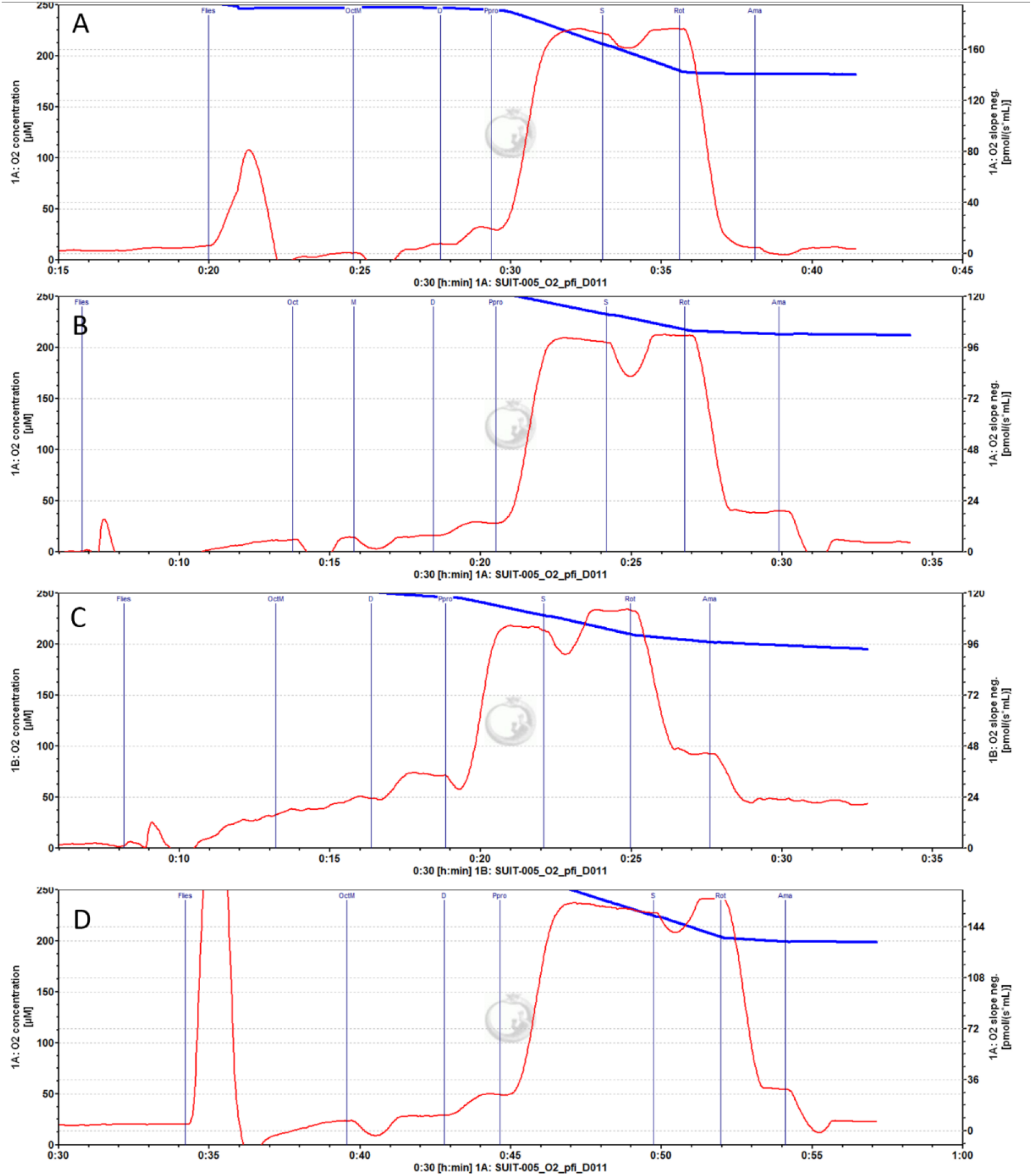
Example Datlab traces from male *D. melanogaster* exposed to **(A)** DMSO, **(B)** 1 mM clozapine, **(C)** 1 mM clozapine + 0.24 mM simvastatin **(D)** 0.24 mM simvastatin for 14 days. Oxygen concentration is shown in blue and oxygen flux in red. Protocol used was SUIT-005 D011 without uncoupler. Marks were made after stabilisation of oxygen flux for a few minutes following reagent administration.

**Supplementary Figure 3:**
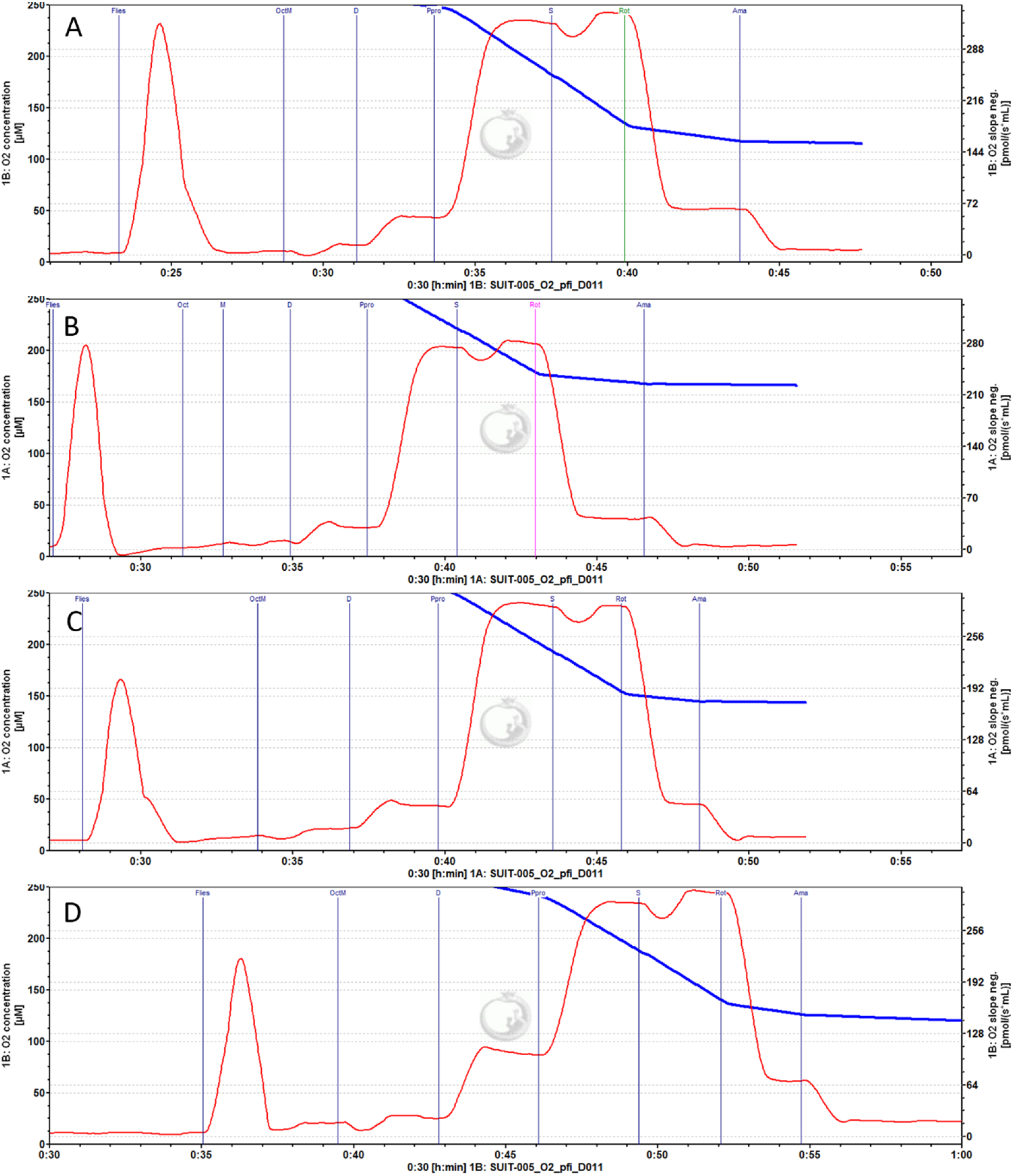
Example Datlab traces from female *D. melanogaster* exposed **(A)** DMSO, **(B)** 1 mM clozapine, **(C)** 1 mM clozapine + 0.24 mM simvastatin **(D)** 0.24 mM simvastatin for 14 days. Oxygen concentration is shown in blue and oxygen flux in red. Protocol used was SUIT-005 D011 without uncoupler. Marks were made after stabilisation of oxygen flux for a few minutes following reagent administration.

